# *Corylus avellana* disease management: using metagenomics to illuminate the rhizosphere microbiome of hazelnut

**DOI:** 10.1101/2025.10.22.684018

**Authors:** John K. Robinson, John M. Steele, Thomas J. Molnar, Sharon Regan, George C. diCenzo

**Author notes:** **Corresponding author:** George C. diCenzo.

## Abstract

The European hazelnut, *Corylus avellana*, is one of the most economically important tree nut crops globally. The biotrophic ascomycete pathogen *Anisogramma anomala*, found naturally associated with wild *C. americana*, continues to pose a significant threat to European hazelnut production across North America. Here, metagenomics was used to examine the taxonomic and functional features of the rhizosphere microbial communities of hazelnut trees differing in their levels of resistance to *A. anomala*: highly tolerant *Corylus americana*, and resistant and susceptible *Corylus avellana*. No statistically significant differences in microbial alpha diversity or beta diversity were noted between the three rhizosphere groups. Compared to bulk soil, all three rhizosphere groups were enriched for the fungal phylum Basidiomycota and bacterial phylum “*Candidatus* Rokubacteriota”. At the genus level, the bacterial genera *Actinospica*, *Occallatibacter*, and “*Candidatus* Sulfotelmatobacter” were under-represented, while the genus *Rhizobacter* was over-represented, in the resistant and susceptible *C. avellana* rhizosphere samples compared to the bulk soil. A total of 45 dereplicated, high-quality metagenome-assembled genomes (MAGs) were generated, corresponding to 41 bacteria and 4 archaea. Many of the MAGs carried multiple biosynthetic gene clusters, including MAGs corresponding to the genera *Lysobacter* and *Actinospica*. Overall, the low differentiation of the rhizosphere microbiomes suggest that differences in *A. anomala* disease expression are likely not associated with differences in the rhizosphere microbiome. Nevertheless, the results shed new light on the rhizosphere communities of two species of hazelnut, and woody perennials more broadly, and identify potential avenues for future research into the development of microbial inoculants for *Corylus* spp..

## INTRODUCTION

Hazelnut cultivation has a rich history and is still of economic importance in contemporary societies. The European hazelnut *Corylus avellana* has been cultivated since Roman times (Boccacci and Botta 2009) and remains one of the world’s principal tree nut crops (Helmstetter et al. 2020). Approximately 70% percent of the world’s hazelnuts are presently grown in Turkey (Food and Agricultural Organization of the United Nations 2021), but the future of Turkey’s hazelnut industry remains uncertain. To stabilize existing hazelnut production and supply growing global demand, the world’s hazelnut orchards must expand across suitable growing regions. Historically, commercial production of *C. avellana* in North America outside of the Pacific Northwest was limited due to the devastating impact of the biotrophic ascomycete pathogen *Anisogramma anomala* (Molnar 2011; Revord et al. 2020; Thompson et al. 1996). Unlike the native but not-commercially amenable American hazelnut species, *Corylus americana*, that is native to North America and resistant or highly tolerant of *A. anomala* infection, European hazelnut is generally highly susceptible. In susceptible hosts, *A. anomala* infection results in the formation of perennial cankers that girdle branches, culminating in tree death (Pinkerton 1995). Current management practices are both costly and labour intensive relying on pruning and large quantities of fungicides (Jeger et al. 2018). Further, it is documented that the *C. avellana* host protection imparted through widely disseminated resistance (R) genes from *C. avellana* ‘Gasaway’ in Oregon and other R genes mapped to the same linkage group, have been overcome in certain regions (Jacobs et al. 2024a). Consequently, there is a pressing need to identify alternate strategies, such as biocontrol, to reduce the impact of *A. anomala* on hazelnut cultivation.

A potential source of biocontrol agents is the native microbial community associated with a plant species of interest. The plant microbiome is typically divided into three major regions: the phyllosphere, representing the above-ground surface regions occupied by microbes; the endosphere, representing the regions comprised of the plant’s inner tissues; and the rhizosphere, which refers to the thin layer of soil surrounding plant roots (Gupta et al. 2021; Santoyo 2022; Zhang et al. 2021). The rhizosphere ecosystem is particularly complex (Jones and Hinsinger 2008) laden with an exceptional diversity of life (Dlamini et al. 2022; Zhang et al. 2020). These plant root-associated microbial communities have been referred to as the plant’s second genome (Berendsen et al. 2012), as they are capable of profoundly extending the plant’s functional potential (Bakker et al. 2013a). Notable examples of plant benefits conferred through these associations include enhanced nutrient availability and improved tolerance to both biotic and abiotic stress (Bakker et al. 2013b; Hakim et al. 2021; Pantigoso et al. 2022).

Importantly, rhizosphere microbial populations can be influenced by both the plant genotype and its disease state (Liu et al. 2019; Liu et al. 2021; Liu et al. 2023a). Though conflicting evidence has been reported (Fitzpatrick et al. 2018), increasing host plant phylogenetic distance has been associated with less similar bacterial rhizosphere composition (Bouffaud et al. 2014; Lei et al. 2019; Liu et al. 2023b), while two genotypes of the same plant species can possess similar but distinguishable microbial populations (Peiffer et al. 2013). Likewise, domestication has been found to influence microbiome assembly and metabolic functions (Yue et al. 2023).

At the same time, rhizosphere microbial communities may be modified based on plant root secretions, with plants potentially recruiting and stimulating disease suppressive microbial taxa in response to pathogens as a means of improving disease outcomes (Liu et al. 2023a). Rhizosphere microbes may mitigate plant disease through antagonistic microbe-microbe interactions (Hassani et al. 2018), priming of the plant immune system via induced systemic resistance (ISR) (Yu et al. 2022), and enhanced nutrient acquisition (Dordas 2008; Dotaniya and Meena 2015). Microbes that exhibit these properties may therefore prove useful in disease management, through serving as biopesticides, enhancing systemic plant disease tolerance through eliciting ISR, or by bolstering overall plant vigour through facilitating nutrient acquisition. The rhizosphere microbiome is therefore worth examining as a promising reservoir of disease attenuating agents, and characterizing interactions between plant roots and their surrounding organisms has demonstrated extensive promise for the improvement of economically valuable crops (Orozco-Mosqueda et al. 2022).

Unfortunately, the rhizosphere of woody perennial plants remains understudied relative to annual crops. Despite the relative paucity of research in this area, the diverse utility of woody perennial associated rhizosphere microbes has been validated by numerous studies. For example, phosphate solubilizing microbes with plant growth promoting potential have been reported in the peach rhizosphere (Li et al. 2025), highlighting the prospect of leveraging rhizosphere microbial communities to enhance woody perennial vigour. In addition, the promise of certain walnut-associated plant growth-promoting rhizobacteria (PGPR) for enhancing drought tolerance (Lotfi et al. 2022) illustrates the abiotic stress ameliorating potential of microbes in the woody perennial rhizosphere. Further, examples of the capacity of rhizosphere microbes to aid in combating biotic stress have been reported for woody perennial crops including apple (Berdeni et al. 2018; Derikvand et al. 2023), pear (Zhang et al. 2023), plum (Ali et al. 2018), and citrus (Ezrari et al. 2021). Collectively, these studies suggest woody perennial rhizosphere microbes may aid in addressing several significant challenges facing woody perennial cultivation.

In the current study, shotgun metagenomics was used to investigate the rhizosphere microbiome of North American grown hazelnut plants. To investigate the effect of genotype and disease resistant phenotypes, samples were collected from an orchard infected with *A. anomala* and were taken from the rhizospheres of native *C. americana* plants and *C. avellana* genotypes that are resistant or susceptible to *A. anomala* infection. By gaining insight into the taxonomic and functional features of hazelnut trees, this study broadens the scope of available genomic information relevant to hazelnut improvement, including for disease management, and may help guide future identification of biocontrol agents.

## METHODS

### Soil sample collection

Bulk soil and rhizosphere samples were collected from Hort Farm 3, 67 Ryders Lane, East Brunswick, NJ 08816, between 1:00 PM and 4:00 PM on 19 October 2022. Soils were collected using a small shovel that was sterilized with 70% ethanol before each sample collection, and transferred to sterile 50 mL tubes. Soils from highly tolerant *C. americana* trees were collected from row 1 of the farm. Soils from *C. avellana* trees collected from row 3 were divided into two groups segregating for disease resistance, designated resistant and susceptible, based on the presence or absence of pronounced *A. anomala* cankers. Five soil samples were collected for each of the three hazelnut varieties; samples were collected at a depth of approximately 15 cm, approximately 15 cm away from the trunk of the tree. In addition, five bulk soil samples were collected at a depth of approximately 15 cm from five sites around the periphery of the hazelnut orchard.

### DNA isolation and sequencing

Approximately one hour after sample collection was complete, ∼0.42 g subsamples of soil were taken from each soil sample. DNA extraction was performed using a Qiagen DNeasy PowerSoil Pro Kit, according to the manufacturer’s instructions. One of the bulk soil samples was omitted from downstream analysis due to failure of the DNA extraction. All other DNA samples were subsequently refrigerated until they were shipped to Génome Québec (Montréal, Quebec, Canada) for library preparation and Illumina sequencing. For samples B2, B3, B5, R1P5, R1P9, R1P14, R3P3, R3P15, R3P20, R3P9-, R3P10-, R3P12-(**Table S1**), library preparation was performed using the NxSeq® AmpFREE Low DNA Library Preparation Kit (Lucigen) using the Xp protocol, as suggested by the manufacturer. These samples were sequenced using 150-bp paired-end technology on an Illumina NovaSeq 6000 instrument. For samples B4, R1P1, R1P18, R3P5, R3P11, R3P14-, R3P18-(**Table S1**), library preparation was performed using the Illumina® DNA PCR-Free Prep, Tagmentation Kit following the manufacturer’s recommendations. These samples were likewise sequenced using 150-bp paired-end technology, but on an Illumina NovaSeq X Plus instrument.

### Taxonomic classification of the raw Illumina reads

Illumina reads were taxonomically classified using Kraken (v2.1.3) (Wood et al. 2019) using a database consisting of the bacteria, archaea, fungi, and virus reference libraries, and post-processed using Bracken (v2.7) (Lu et al. 2017). Alpha diversity and beta diversity measurements were determined based on the species-level output of Bracken. Diversity measurements were determined, analyzed, and visualized in R (v4.5.1) (R Core Team 2025) using the tidyverse (v2.0.0) (Wickham et al. 2019), dplyr (v1.1.4) (Wickham et al. 2018), phyloseq (v1.52.0) (McMurdie and Holmes 2013), stats (v4.5.1) (R Core Team 2025), ggpubr (v0.6.1) (Kassambara 2025), vegan (v2.7.1) (Oksanen et al. 2024), pairwiseAdonis (v0.4.1) (Martinez Arbizu 2017), ggplot2 (v4.0.0) (Wickham 2009), and patchwork (v1.3.2) (Pedersen 2019) packages. KrakenTools (v1.2) (Lu et al. 2022) and KronaTools (v2.8) (Ondov et al. 2011) were used to generate krona plots.

### Metagenome assembly and binning

Illumina reads were filtered using BBduk (v39.06) (Bushnell 2014) and trimmed using Trimmomatic (v0.39) (Bolger et al. 2014) with the parameters: LEADING:3 TRAILING:3 SLIDINGWINDOW:4:15 MINLEN:36. Following trimming, metagenomes were assembled using MEGAHIT (v1.2.9) (Li et al. 2015) with the “presets” option set to “meta-large”. Metagenome assembly statistics were calculated using the stats.sh script of BBMap (v37.36) (Bushnell 2014). In addition, to confirm that the alpha and beta diversity metrics were not confounded by differences in sequencing depth, seqtk (v1.3) (https://github.com/lh3/seqtk) was used to filter the trimmed reads to obtain a subset corresponding to 90% of the number of reads from the sample with the fewest reads. These reduced read sets were then used for metagenome assembly using MEGAHIT with the “presets” option set to “meta-large”.

### Assembly of metagenome-assembled genomes

Prior to binning, contigs less than 1500 bp were discarded using pullseq (v1.0.2) (https://github.com/bcthomas/pullseq). Next, an all-against-all mapping of the Illumina reads against each metagenome was performed using bowtie2 (v2.4.5) (Langmead and Salzberg 2012) and sorted using samtools (v1.15.1) (Li et al. 2009). Binning was then performed independently for each sample using MetaBAT2 (v2.14) (Kang et al. 2019), CONCOCT (v1.1.0) (Alneberg et al. 2014), MaxBin2 (v2.2.7) (Wu et al. 2016), and SemiBin2 (v2.1.0) (Pan et al. 2023). From each of the metagenomes, an optimized set of MAGs was produced using DAS_Tool (v1.1.4) (Sieber et al. 2018) and the output of the four binning algorithms. In addition, bins were generated with VAMB (v4.1.3) (Nissen et al. 2021, 20), using all metagenomes and with read mapping performed using minimap2 (v2.24-r1122) (Li 2018). Finally, a non-redundant set of high-quality MAGs (> 70% completeness; < 10% contamination) was generated from all the produced MAGs using dRep (v3.4.1) (Olm et al. 2017) and a secondary clustering threshold of 99%. The dependencies for dRep included Mash (v2.3) (Ondov et al. 2016) MUMmer (v4.0.0rc1) (Kurtz et al. 2004), CheckM (v1.2.2) (Parks et al. 2015), Prodigal (v2.6.3) (Hyatt et al. 2010), and pplacer (v1.1.alpha19) (Matsen et al. 2010).

### Taxonomic classification of the assembled metagenomes

Prior to taxonomic classification, contigs less than 500 bp were discarded using pullseq. Contigs were taxonomically classified using MMSeqs2 (v0a6b8c) (Steinegger and Söding 2017) with the UniRef90 database (downloaded 18 July 2024). Custom scripts were used to convert the MMSeqs2 output data into taxa abundance tables, with taxa abundances calculated as the sum of contig length multiplied by contig sequencing depth for all contigs with a given taxonomic classification. Alpha diversity and beta diversity metrics were determined, analyzed, and visualized in R using the tidyverse, ggplot2, dplyr, phyloseq, stats, ggpubr, vegan, pairwiseAdonis, and patchwork packages. The relative abundances of the taxa with an average relative abundance of at least 1.0% were visualized as bubble plots and stacked bar plots with R’s ggplot2, patchwork, and randomcoloR (v1.1.0.1) (Ammar 2019) packages. Lastly, the R packages ANCOMBC2 (v2.10.1) (Lin and Peddada 2024) and microbiome (v1.30.0) (Lahti and Shetty 2019) were used to identify statistically significant differences in taxa abundances for taxa with an average relative abundance of at least 0.1% across samples. Differentially abundant taxa were then visualized in R using the pheatmap (v1.0.13) (Kolde 2025) package.

### Taxonomic classification of metagenome-assembled genomes

MAGs were taxonomically classified using GTDB-Tk (v2.4.0) with the R220 database and the classify_wf workflow (Chaumeil et al. 2020). To construct a phylogeny of the bacterial MAGs, the identify function of GTDB-Tk was used to identify homologs of the bac120 gene set, following which the align function of GTDB-Tk was used to align each marker set and then concatenate the alignments. Lastly, the concatenated alignment was used to construct a maximum likelihood phylogeny using IQ-TREE2 (v2.2.2.4) (Minh et al. 2020). First, ModelFinder (Kalyaanamoorthy et al. 2017), was used to identify the best scoring model based on Bayesian information criterion (BIC). Then, IQ-TREE2 was run using the best-scoring model (LG+F+R5) with branch support assessed using the Shimodaira–Hasegawa-like approximate likelihood ratio test (SH-aLRT) and ultrafast jackknife analysis with a subsampling proportion of 40%, with both metrics calculated from 1,000 replicates. Phylogenies were visualized using iTOL (Letunic and Bork 2024).

### Characterization of the metabolic potential of the metagenomes and MAGs

Prior to functional annotation, contigs less than 1000 bp were discarded from the metagenomes using pullseq. Next, DRAM (v0.1.2) (Shaffer et al. 2020), as implemented on the KBase platform (Arkin et al. 2018), was used to annotate all metagenomes and MAGs. For the metagenomes, the KEGG annotations were extracted from the annotation file, and a Bray-Curtis distance matrix was produced and visualized using a Principal Coordinate Analysis in R with the ggplot2, dplyr, phyloseq, and patchwork packages. Statistical analyses were performed in R using the vegan and pairwiseAdonis packages. Group specific functional differences were determined and visualized using the dplyr, stats, dunn.test (v1.3.6) (Dinno 2024), and ggplot2 packages.

Prior to annotation of BGCs, contigs less than 500 bp were discarded from the metagenomes using pullseq. BGCs were identified in the reduced metagenomes using antiSMASH (v7.0) (Blin et al. 2023) with default parameters, after which BiG-SCAPE (v1.1.8) (Navarro-Muñoz et al. 2020) was run with default parameters to examine BGC diversity across the metagenomes. Likewise, BGCs were identified in the MAGs using antiSMASH and BiG-SCAPE. The output files were processed and visualized as heatmaps in R using the dplyr, ggplot2 and patchwork packages.

## RESULTS

### Hazelnut rhizosphere sampling

Rhizosphere samples were collected from 15 hazelnut trees situated at Rutgers University Hort Farm 3, East Brunswick, New Jersey, USA, in October 2022. Five samples were collected from each of the following tree types: native *C. americana* plants (henceforth referred to as “native”), *C. avellana* resistant to *A. anomala* (henceforth referred to as “resistant), and susceptible *C. avellana* (henceforth referred to as “susceptible”). The *C. americana* plants were derived from open pollinated seeds obtained from wild plants in Dearborn, Michigan, whereas the *C. avellana* plants were the product of a controlled cross of a H4AR21P05 (Jacobs et al. 2024b) female (open pollinated seed from Warsaw, Poland) with a OSU 474.084 (‘Lewis’ x ‘Tonda di Giffoni’) x OSU 540.084 (a full sibling of ‘Sacajawea’) male pollen source, obtained from Shawn Mehlenbacher of Oregon State University. These *C. avellana* plants were expected to segregate for a resistance (R) gene originating from Poland (Capik et al., 2013). The two groups of *C. avellana* trees used for this study, therefore, represent full siblings with the resistant group expressing no signs or symptoms of disease and the susceptible expressing obvious signs of infection with pronounced cankers and stem dieback predicated on segregation of the major R gene. At the time of collection, the trees were ∼6 years of age and have maintained consistent disease phenotypes for ∼8 years as of 2024. All rhizosphere samples were collected from soils adjacent to tree roots, at a depth of ∼ 15 cm and a radius of ∼ 15 cm from the trunk of the tree. For comparison, five bulk soil samples (not near tree roots) were also collected from a depth of ∼ 15 cm around the orchard; however, the DNA isolation from one bulk soil sample failed, leaving us with four bulk soil samples.

### Metagenomic sequencing and assembly statistics

Illumina 150 bp paired-end sequencing was used to characterize the microbial community of each of the soil samples, generating between ∼ 35 million and ∼118 million paired-end reads per sample (**Table S1**), of which between ∼ 31 million and ∼ 113 million survived quality control (**Table S1**). Metagenome assembly using MEGAHIT resulted in assemblies of 544 Mb to 4007 Mb in size, with N50 values between 446 kb and 2260 kb (**Table S1**). When limiting the metagenomes to contigs of at least 0.5 kb in size, the assemblies averaged 612 Mb and contained an average of 768,075 contigs (**Table S1**). Likewise, when limiting the metagenomes to contigs of at least 1 kb in size, the assemblies averaged 187 Mb and contained an average of 115,156 contigs (**Table S1**).

### Taxonomic characterization of the hazelnut rhizosphere

Initially, the community composition of each rhizosphere and bulk soil sample was evaluated based on the raw Illumina reads using Kraken2 and Bracken, with a database consisting of the bacteria, archaea, fungi, and virus reference libraries. However, the rate of classification by Kraken2 was low, with just 0.80% to 1.77% of reads assigned a taxonomy (**Table S2**). Consequently, we instead used MMSeqs2 with the UniRef90 database, to taxonomically classify the assembled contigs (minimum contig size of 0.5 kb); using this approach, contigs encompassing between 6% and 31% of reads were assigned a taxonomy (**Table S2**). Overall, the metagenomes were dominated by bacteria, with 82% to 96% of reads mapping to the classified contigs corresponding to bacteria, compared to 1.0% to 9.5% and 0.3% to 3.8% for archaea and eukaryotes, respectively.

Alpha diversity (Shannon diversity index and Simpson diversity index [1-D]) was calculated from the MMSeqs2 output (**Figures 1A, 1B**). No statistically significant differences were observed between sample groups using either of the diversity measures. However, both diversity metrics displayed a trend of the bulk soil samples having lower diversity than the rhizosphere samples; indeed, statistically significant differences were detected for both diversity metrics when comparing bulk soil versus all rhizosphere samples grouped as a single treatment (Kruskal-Wallis p-value < 0.05 for both metrics). In addition, although not statistically significant, there appeared to be greater between-replicate variance in the diversity metrics for the non-native *C. avellana* compared to native *C. americana* samples, perhaps suggesting that the rhizosphere microbial community of the non-native plants was less stably assembled than that of the native plants.

**Figure 1.**
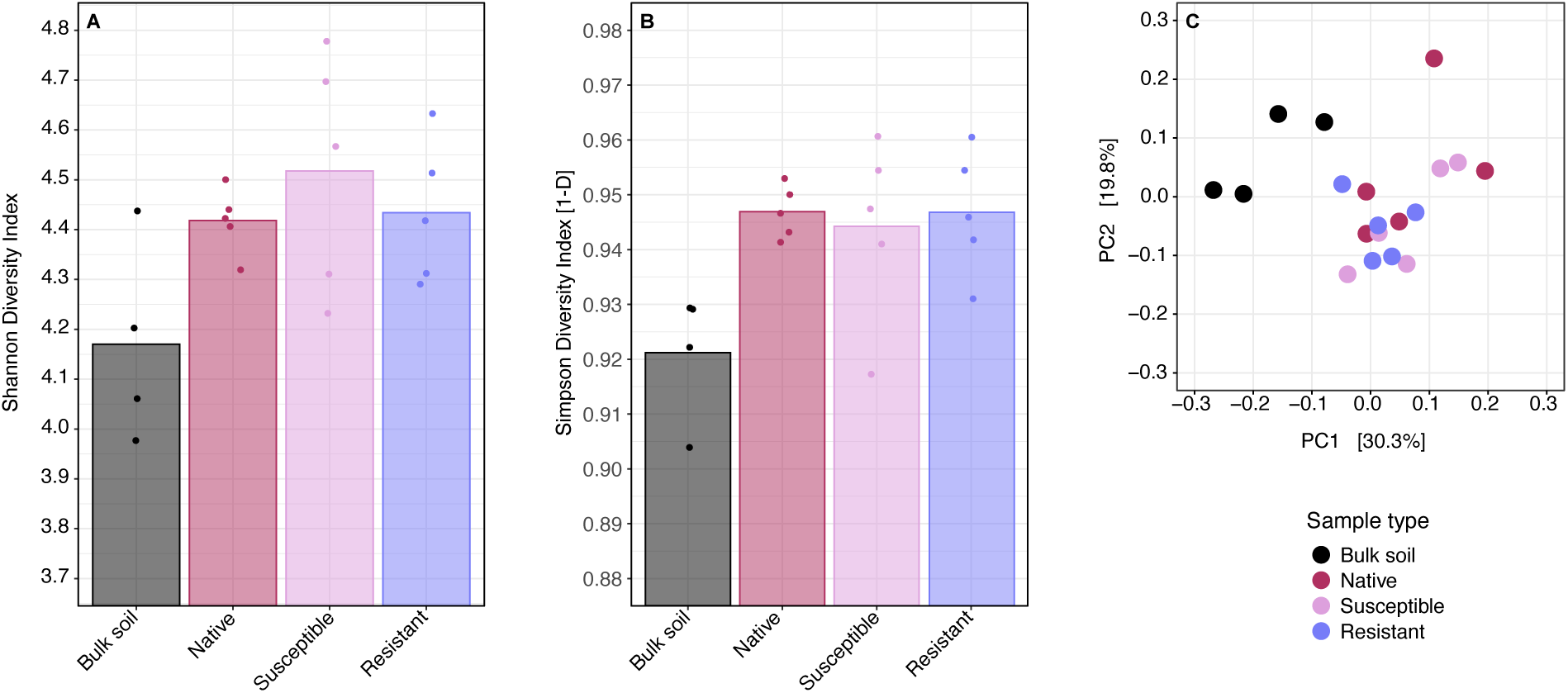
Diversity metrics of the metagenomes. (**A**, **B**) Alpha diversity values calculated using the (A) Shannon Diversity Index (Kruskal-Wallis rank sum test, p-value = 0.16) or (B) Simpson Diversity Index [1-D] (Kruskal-Wallis rank sum test, p-value = 0.06) are shown. Alpha diversity was calculated based on species level taxonomic classification of assembled contigs (minimum length of 500 bp). (**C**) A Principal Coordinate Analysis (PCoA) plot based on Bray-Curtis dissimilarity values is shown (PERMANOVA, p-value < 0.001), with pairwise adonis tests indicating that the bulk soil was statistically different from the rhizospheres of the native (p.adjusted = 0.048), susceptible (p.adjusted = 0.047), and resistant (p.adjusted = 0.047) varieties.

Beta diversity was calculated at the species level using Bray-Curtis dissimilarity (**Figure 1C**). Statistically significant differences (PERMANOVA p-value < 0.001) in beta diversity were observed between the control and each of the three rhizosphere sample groups, providing support that the rhizosphere samples were correctly collected and were reflective of the rhizosphere community. However, no significant differences were observed between the native, resistant, and susceptible rhizosphere samples, suggesting that plant genotype or disease state had little to no impact on the rhizosphere microbiome at a global level.

To ensure that the diversity metrics were not being skewed by differences in read depth, the sequencing reads were subsampled to achieve equivalent read numbers (following QC), after which metagenomes were reassembled and MMSeqs2 rerun. The alpha and beta diversity metrics were then recalculated from these data. This process produced qualitatively similar results for the alpha and beta diversity metrics (**Figure S1**), indicating that the observed diversity values were not heavily skewed by differences in read counts between samples.

### Differentially abundant taxa

The relative abundances of major taxa were investigated across samples at the phylum and genus levels (**Figures 2 and 3**). Initial, qualitative examination of the data revealed that all samples were dominated by the bacterial phyla *Actinomycetota*, *Pseudomonadota*, and *Acidobacteriota* (**Figure 2A**). At the genus level, the three most abundant taxa were the common soil bacteria *Bradyrhizobium*, *Gaiella*, and *Nocardioides* (**Figure 3A**). Although the archaea represented only a minority of the metagenomes, the archaeal component was dominated by the phylum *Thermoproteota* (syn. *Nitrosophaerota*) (**Figure 2B**), driven by the abundance of the genera *Nitrososphaera* and “*Candidatus* Nitrosocosmicus” (**Figure 3B**). Lastly, examination of the eukaryotic taxa revealed the phylum Basidiomycota was substantially enriched in the rhizosphere samples relative to the bulk soil control, while the phylum Ascomycota also appeared somewhat enriched in many of the rhizosphere samples (**Figure 2C**). The differential abundance of the phylum Basidiomycota was driven mostly by the genera *Hebeloma*, *Laccaria*, *Galerina*, and *Scleroderma*, while the differential abundance of the phylum Ascomycota was driven mostly by the genus *Morchella* and to a lesser extent the genus *Terfezia* (**Figure 3C**). It is important to note, however, that some unexpected eukaryotic taxa were identified in relatively high abundance, such as the shark genus *Chiloscyllium* (**Figure 3C**). This may be the result of poor taxonomic classification of the eukaryotic contigs and/or the overall low abundance of eukaryotes in the data. Regardless, it does suggest that caution should be taken in interpreting the abundance data for the eukaryotic taxa.

**Figure 2.**
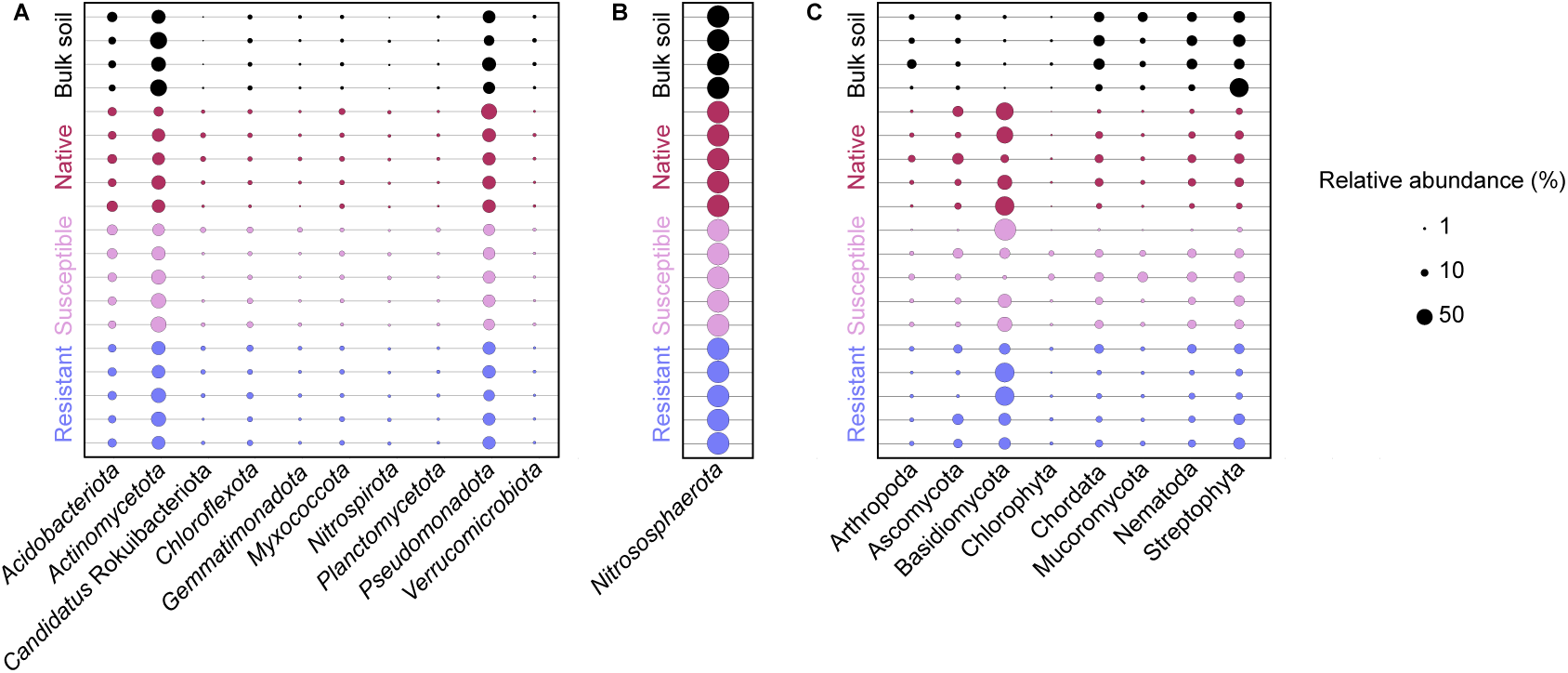
Relative abundance of phyla across the metagenomes. The abundance of (**A**) bacterial phyla, (**B**) archaeal phyla, and (**C**) eukaryotic phyla, displayed as a percentage of the total bacteria, archaea, and eukaryotes, respectively. The rows represent each of the metagenomic samples, colour coded by group, while the columns represent taxa with an average relative abundance of at least 1% of the bacteria, archaea, or eukaryotes, respectively. The size of the dots corresponds is proportional to the relative abundance of the taxon in a given sample, as indicated by the legend to the right of the figure.

**Figure 3.**
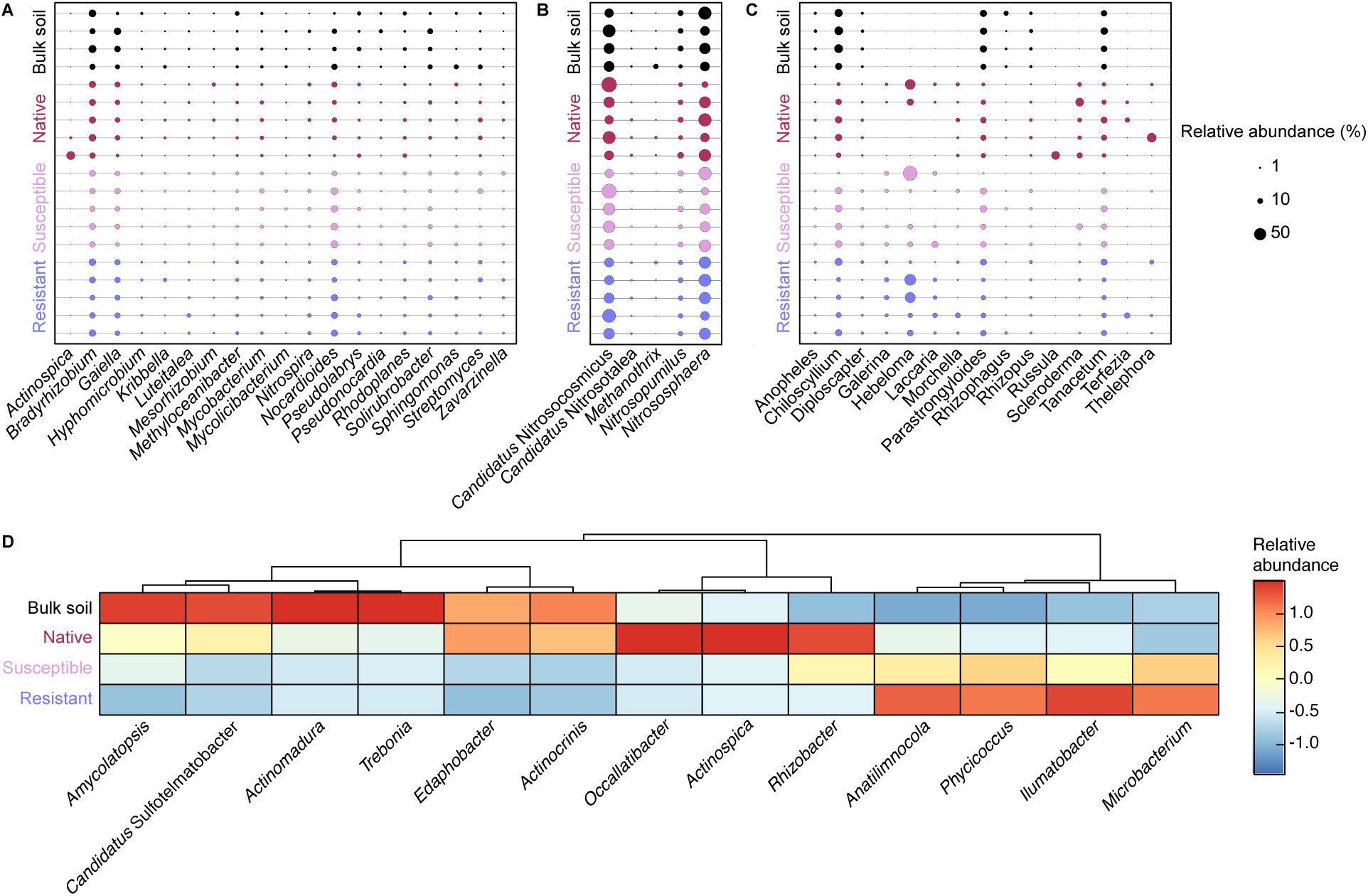
Relative abundance of genera across the metagenomes. The abundance of (**A**) bacterial genera, (**B**) archaeal genera, and (**C**) eukaryotic genera, displayed as a percentage of the total bacteria, archaea, and eukaryotes, respectively. The rows represent each of the metagenomic samples, colour coded by group, while the columns represent taxa with an average relative abundance of at least 1% of the bacteria, archaea, or eukaryotes, respectively. The size of the dots corresponds is proportional to the relative abundance of the taxon in a given sample, as indicated by the legend to the right of the figure. (**D**) A heatmap showing the differential abundances of bacterial genera identified as showing statistically significant differences in abundances across treatments. Columns were scaled using pheatmap in R. Dark red indicates higher abundances while dark blue indicates lower abundances.

Following qualitative assessment of the abundance data, ANCOMBC2 was used for quantitative assessment of differential abundances of the taxa accounting for at least 0.1% of the overall relative abundances of the communities. At the phylum level, the eukaryotic phylum Basidiomycota and the bacterial phylum “*Candidatus* Rokuibacteriota” appeared enriched in all three rhizosphere groups relative to the bulk soil controls with (log2[fold change] ≥ 3.9 and 2.3, for Basidiomycota and “*Candidatus* Rokuibacteriota”, respectively; **Figure 2A**). On the other hand, the bacterial phylum *Verrucomicrobiota* appeared underrepresented specifically in the rhizosphere communities of the “resistant” *C. avellana* plants relative to the bulk soil controls (log2[fold change] of -1.2; **Figure 2A**). At the genus level, the three rhizosphere groups were depleted in the bacterial genera *Trebonia* (log2[fold change] ≤ -2.6) and *Actinomadura* (log2[fold change] ≤ -1.3) relative to the bulk soil (**Figure 3D**). Compared to the bulk soil controls, the resistant and susceptible *C. avellana* samples (but not the *C. americana* samples) both had an underrepresentation of the bacterial genera *Actinospica* (log2[fold change] ≤ -1.4), *Occallatibacter* (log2[fold change] ≤ -2.8) and “*Candidatus* Sulfotelmatobacter” (log2[fold change] ≤ -2.9), while the bacterial genus *Rhizobacter* appeared enriched (log2[fold change] ≥ 1.5) (**Figure 3D**). The resistant *C. avellana* samples also appeared depleted in the bacterial genera *Amycolatopsis* (log2[fold change] of -1.2), *Actinocrinis* (log2[fold change] of -1.4), and *Edaphobacter* (log2[fold change] of -1.7) and enriched in *Phycicoccus* (log2[fold change] of 1.3), *Ilumatobacter* (log2[fold change] of 1.9), and *Anatilimnocola* (log2[fold change] of 1.8) relative to the bulk soil (**Figure 3D**). Lastly, the bacterial genus *Microbacterium* was enriched in the resistant *C. avellana* samples relative to the native variety (log2[fold change] of 2.0) (**Figure 3D**). *Microbacterium* also visually appeared to be enriched in the susceptible *C. avellana* compared to the native variety although this difference was not statistically significant.

### Functional overview of the metagenomes

DRAM was used for functional annotation of all contigs of at least 1 kb in length, and the relative abundance of genes assigned KEGG annotations were used to construct a PCoA plot using Bray-Curtis dissimilarities as a general overview of the functional differentiation of each sample (**Figure 4**). Although global differences between samples were detected (PERMANOVA p-value of 0.031), no statistically significant differences were noted for any of the pairwise comparisons between sample groups. However, although not statistically significant, there did appear to be a slight trend towards the native *C. americana* and resistant *C. avellana* samples grouping together and separating from the sensitive *C. avellana* and bulk soil. Likewise, no clear differences were noted between sample types when comparing the presence of various functions and pathways related to carbohydrate metabolism, methanogenesis and methanotrophy, nitrogen metabolism, short-chain fatty acid and alcohol conversions, and sulfur metabolism (**Figure S2**). Overall, these results indicate that there was little functional differentiation between the sample types at a global level.

**Figure 4.**
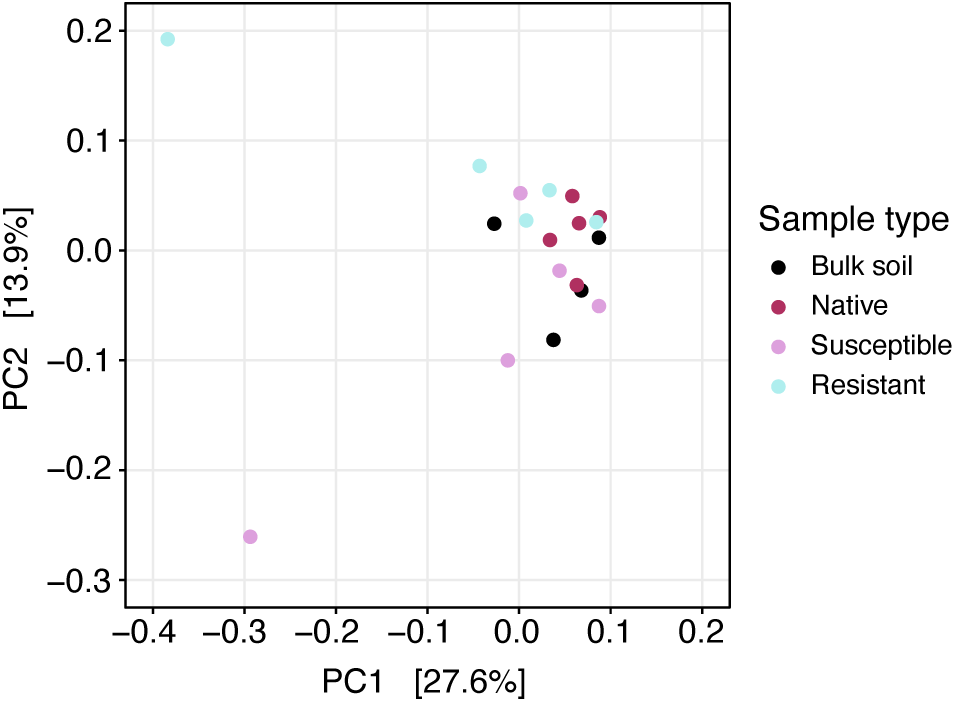
Global comparison of the metabolic potential of the metagenomes. A Principal Coordinate Analysis (PCoA) plot based on Bray-Curtis dissimilarity values calculated from the KEGG annotations produced for each metagenome using DRAM. While statistically significant global differences were observed (PERMANOVA, p-value = 0.031), no statistically significant differences between groups were detected using pairwise adonis tests.

In addition, prokaryotic biosynthetic gene clusters (BGCs) were annotated in all metagenome assemblies (limited to contigs of at least 0.5 kb) using the program antiSMASH. BiG-SCAPE was then used to group the antiSMASH annotations based on the class and predicted products of the BGCs (**Figure 5**). The distribution of BGCs across samples was largely similar, regardless of the sample group, with BGCs associated with aryl polyenes, indoles, nonribosomal peptides, phosphonates, ribosomally synthesized and post-translationally modified peptides, polyketides, terpenes, and thioamitides detected across the majority of samples.

**Figure 5.**
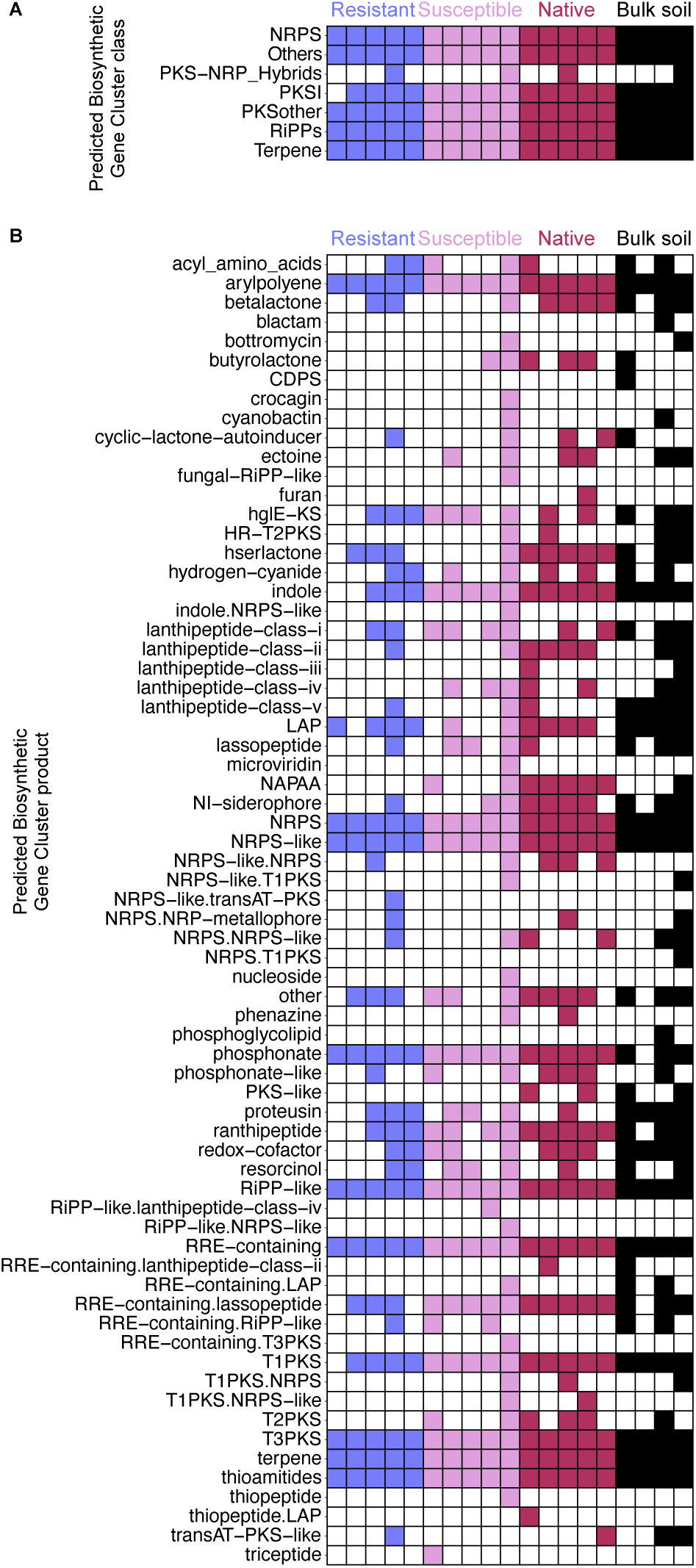
Putative biosynthetic gene clusters detected in the metagenomes. (**A**) A heatmap showing distribution of predicted biosynthetic gene clusters summarized by biosynthetic gene cluster classification. (**B**) A heatmap showing the distribution of predicted biosynthetic gene clusters summarized by predicted product. (**A**, **B**) A coloured box indicates that a biosynthetic gene cluster classification or predicted product was detected in a given metagenome, whereas a white box indicates that the pathway was not detected.

### Reconstruction and characterization of metagenome-assembled genomes

Following dereplication using an average nucleotide identity (ANI) threshold of 99%, 45 high-quality (completeness ≥ 70%, contamination ≤ 10%) metagenome-assembled genomes (MAGs) were constructed from the dataset and taxonomically categorized with GTDB-Tk, including 41 bacterial and 4 archaeal MAGs (**Figure 6, Table S3**). Interestingly, only one of the 45 MAGs could be taxonomically classified at the species level when using the Genome Taxonomy Database Toolkit (GTDB-Tk) with database R220, while an additional nine were classified to the genus level. The four archaeal MAGs represent four distinct species from three genera within the family *Nitrososphaeraceae* (phylum *Thermoproteota*). On the other hand, nine phyla are represented amongst the 41 bacterial MAGs, with the majority of MAGs belonging to the phyla *Actinomycetota* (21 MAGs), *Pseudomonadota* (9 MAGs), and *Acidobacteriota* (6 MAGs). MAGs were functionally annotated with DRAM, and a summary of the results are provided as **Figure S3**. In addition, antiSMASH and BiG-SCAPE were used to detect and summarize the abundance of BGCs in the MAGs, and these results are summarized in **Figure S4**.

**Figure 6.**
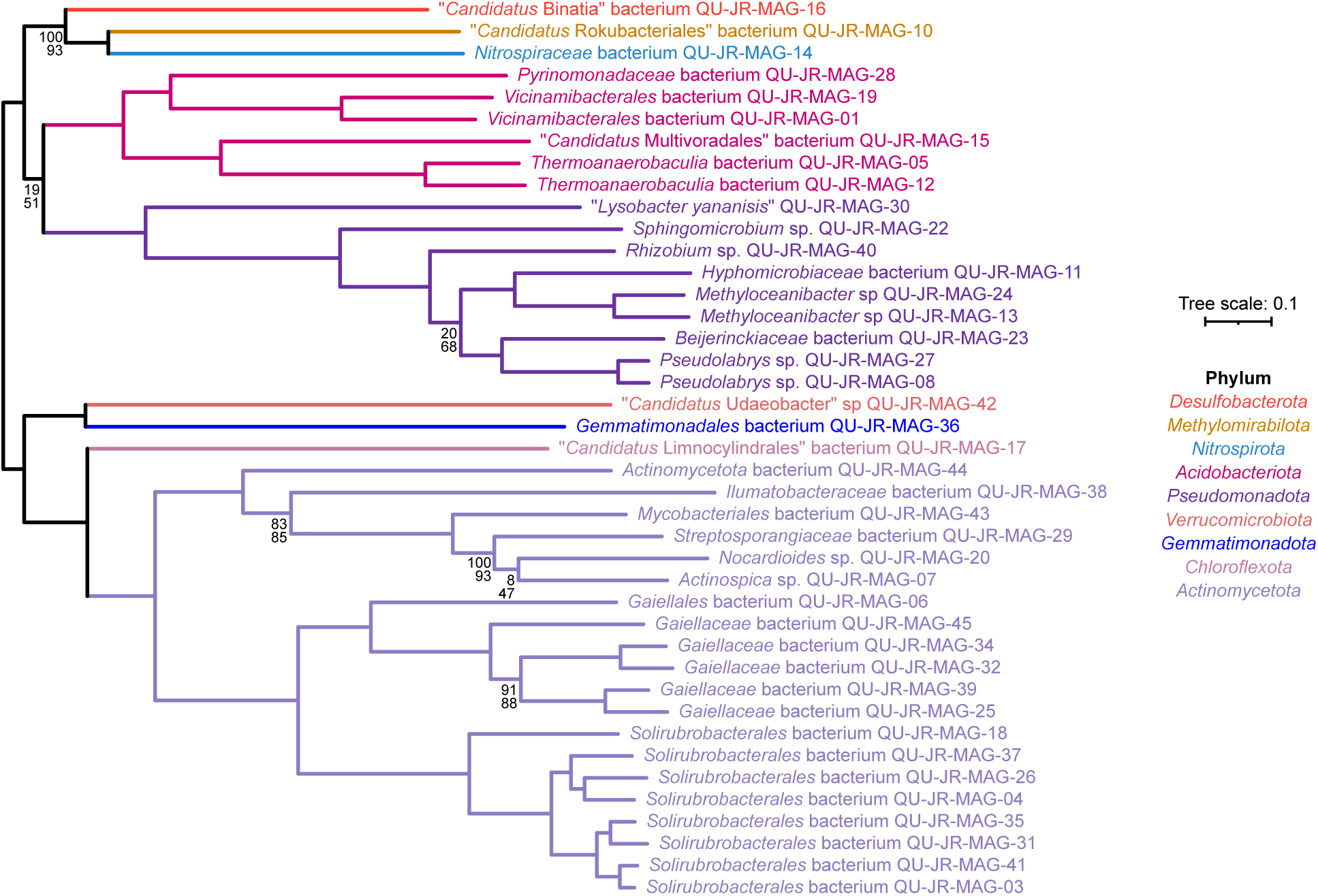
Unrooted phylogeny of the bacterial MAGs generated in this study. An unrooted, maximum likelihood phylogeny of the 41 high-quality metagenome-assembled genomes (MAGs) generated in this study, prepared from a concatenated alignment of the bac120 gene set of the Genome Taxonomy Database. Taxa names and branches are colour coded based on phylum. Thescale represents the mean number of nucleotide substitutions per site. The numbers on the nodes indicate the SH-aLRT support values (top numbers) and the ultra-fast jackknife values using a 40% resampling rate (bottom numbers), both calculated from 1000 replicates; values are only shown at nodes where at least one value is below 95.

## DISCUSSION

### The hazelnut rhizosphere microbiome showed only minor differences compared to bulk soil

Previous studies examining bacterial community structure in the roots of woody plants (*Quercus*, *Populus*, and *Pinus*) found that bacterial community structure was shaped more strongly by host tree species than by soil origin (Bonito et al. 2014). We therefore expected to see differences between the microbial community of the rhizosphere samples compared to the bulk soil samples. As expected, the bulk soil samples were clearly delineated from the rhizosphere samples based on the alpha and beta diversity measures. This differentiation appeared to be mainly driven by changes in the frequencies of low abundance taxa as we observed few differences in the relative abundances of the primary taxa (those with relative abundances ≥ 0.1%). However, the current study examined only a single time point. Considering that another study observed that the prokaryotic component of the maize rhizosphere only differed from bulk soil at specific stages of plant development (Wattenburger et al. 2019), it may be that sampling at a different time of year, or sampling deeper roots, would reveal more dramatic changes in the hazelnut rhizosphere microbial community structure than observed here.

Few differences were observed between the microbial communities of the different hazelnut genotypes, contrary to our expectation (Lopes et al. 2023). Likewise, we observed little difference between *C. avellana* plants that were or were not infected with *A. anomala*, despite previous studies observing that foliar pathogens can indirectly impact rhizosphere microbial communities in ways that could potentially have positive impacts on plant disease tolerance (Berendsen et al. 2018; Jiang et al. 2023; Luo et al., 2022; Yang et al. 2020). While these results could suggest that hazelnut genotype and disease status have little impact on the rhizosphere microbiome, it may also be that differences would be more pronounced at other times of the year, or that deeper sequencing and/or taxonomic classification of a higher portion of the contigs would reveal additional differences missed here.

### Composition of the hazelnut microbial community

We found that the fungal phylum Basidiomycota was enriched in all rhizosphere samples relative to bulk soil. Likewise, the fungal phyla Ascomycota appeared slightly enriched in the rhizosphere samples, although Ascomycota did not meet the 0.1% relative abundance threshold used to retain abundant taxa from the results of ANCOMBC2. Interestingly, a separate study reported Ascomycota, Basidiomycota, and Mortierellomycota as the dominant fungal phyla in the rhizosphere of hazelnut varieties grown in Beijing, China (Ma et al. 2021). Similarly, the phyla Basidiomycota and Ascomycota were the dominant fungal phyla in the rhizosphere of the deciduous shrub *Paeonia ludlowii*, with Basidiomycota being the most dominant in the autumn (Qiao et al. 2023). Overall, the consistency in these results provides support that our rhizosphere samples accurately capture the hazelnut rhizosphere microbiome.

Although there is limited data on the effects of bacteria on hazelnut tree health and growth, one study reported that inoculation of *C. avellana* seedlings with the bacteria *Pseudomonas putida*, *B. subtilis*, and *Enterobacter cloacae* can improve growth (Rostamikia et al. 2017). Therefore, we wished to use our metagenomics data to help identify microbes in the hazelnut rhizosphere microbiome that could potentially be targets to develop into biocontrol agents to limit the impact of *A. anomala* infection. To this end, we focused on the microbes that were either enriched or depleted in the hazelnut microbiome. One taxon of particular interest based on the relative and differential abundance data was the bacterial phylum “*Candidatus* Rokuibacteriota”, which was enriched in all three rhizosphere groups relative to bulk soil. “*Candidatus* Rokuibacteriota” have large genomes with high GC content, with the capacity for multifaceted mixotrophic metabolism (Becraft et al. 2017). Bacteria from this phylum have also been reported to encode numerous biosynthetic loci in their genomes (Sharrar et al. 2020). Consequently, members of this phylum exhibit characteristics of relevance for biocontrol applications, and we suggest that their utility should be explored within this context.

On the other hand, the bacterial genera *Actinospica*, *Occallatibacter*, and “*Candidatus* Sulfotelmatobacter” were depleted in the susceptible and tolerant varieties of *C. avellana* relative to bulk soil, unlike the native *C. americana* variety. *Actinospica* may be of particular note for *A. anomala* disease management, as there is an ongoing interest in the efficacy of acidophilic actinomycetes as biocontrol agents against phytopathogenic fungi (Poomthongdee et al. 2015). Supplementing the abundance of this genus in the *C. avellana* microbiome may therefore be a strategy worth exploring for *A. anomala* disease management.

In terms of eukaryotic microbes, associations with mycorrhizal fungi have been shown to enhance disease tolerance in woody perennials (Dong et al. 2021; Kebert et al. 2022), and a variety of additional benefits have been described for mycorrhizal associated hazelnut trees. In *C. avellana*, association with the ectomycorrhizal forming fungus *Tuber melanosporum* was reported to regulate important plant defense genes, which may favourably impact the host plant’s response to stress (Sillo et al. 2022). Inoculation of hybrid hazel (*C. heterophylla* x *C. avellana*) seedlings with the ectomycorrhizal fungus *Scleroderma bovista* was reported to promote growth, enhancing photosynthetic efficiency and total plant weight (Cheng et al. 2024). In addition, inoculation with the arbuscular mycorrhizal fungus *Glomus iranicum* var. *tenuihypharum* was found to increase the number of female flowers and overall yield in the Tonda di Giffoni variety of *C. avellana* by 60% and 38%, respectively, relative to the uninoculated control (Luciani et al. 2019). In our study, the fungal genera *Hebeloma*, *Galerina*, *Laccaria*, *Morchella*, *Terfezia*, and *Scleroderma* appeared enriched in the three groups of rhizosphere samples, relative to the control group; however, as the overall relative abundance of fungi across samples was generally low, most of these taxa did not meet the 0.1% relative abundance threshold required to be retained in the results from ANCOMBC2. Isolates of species from these fungal taxa from the hazelnut rhizosphere may therefore represent interesting candidates to explore for plant growth promotion and/or biocontrol against *A. anomala* in North America.

### Primary metabolism of the hazelnut rhizosphere microbiome

Rhizosphere microbes play important roles in nutrient cycling and availability (Thepbandit and Athinuwat 2024) and consequently, they can impact plant nutrition and thus disease tolerance. For example, nitrogen availability can impact both physical and biochemical defence mechanisms in plants, yet the consequences of enhanced endogenous nitrogen can be difficult to anticipate, as they may be favourable or detrimental to plant pathogen tolerance depending on context (Sun et al. 2020). In our study, global differences in the KEGG metabolic annotations (PERMANOVA p-value of 0.031) were observed between samples, however no pairwise differences were detected between sample groups.

Pathways involved in nitrogen metabolism were well represented across most samples including ammonia oxidation, conversion of nitrate to nitrite, nitrite to nitrate, nitrite to nitric oxide, and nitric oxide to nitrous oxide. Though less common, some bulk soil and rhizosphere samples possessed the capacity for nitrite reduction to ammonia. As nitrate serves as the major source of nitrogen for most land plants in aerobic soils (Ho and Tsay 2010), the presence of pathways involved in the production of nitrate across samples highlights the capacity of microbes within these systems to facilitate the production of bioavailable nitrogen for hazelnut. Bioavailable sulfur can also have drastic impacts on plant disease outcomes. For example, inorganic sulfate salts can significantly enhance the resistance of plants to certain fungal pathogens, through a process deemed sulfur-induced resistance (SIR) (Künstler et al. 2020). Here, pathways responsible for the conversion of thiosulfate to sulfate were well represented across all samples besides one of the Susceptible samples. As both bioavailable nitrogenous and sulfur-containing compounds have been demonstrated to alter plant disease tolerance (Bloem et al. 2015; Mur et al. 2017) the application of certain microbial communities that alter the abundance of bioavailable forms of these compounds within the soil may be explored as an alternative means of broadly improving plant health.

Many of the MAGs generated in this study, particularly those within the orders *Gaiellales* and *Hyphomicrobiales* (syn. *Rhizobiales*), encoded pathways involved in the conversion of nitrite to nitric oxide (NO). NO can be generated endogenously in plants and performs important functions in various physiological responses and developmental stages (Hancock 2020; Sun et al. 2021), with roles in plant tolerance to abiotic and biotic stress (Fancy et al. 2017, 201; Mur et al. 2017). Exogenous application of NO, through the use of NO donors, has been shown to improve plant disease tolerance in several instances (Nasir et al. 2020; Khan et al. 2021; Bakhoum et al. 2023). Furthermore, studies suggest that leveraging NO-producing organisms to enhance disease tolerance in plants is a promising avenue to explore. For example, a study examining tomato infected with *Ralstonia solanacearum* found that *Pseudomonas fluorescens* strains overproducing NO displayed better biocontrol efficacy than those that do not overproduce NO (Wang et al. 2005). Consequently, it would be interesting to isolate diverse strains from promising genera within the *Gaiellales* and *Hyphomicrobiales* orders, which we found commonly encode the pathways for NO production, and assess them for their impact on hazelnut disease tolerance. Of particularly interest would be isolates that additionally have the potential to convert thiosulfate to sulfate.

### Secondary metabolism of the hazelnut rhizosphere microbiome

Soil microbes represent a rich source of secondary metabolites (Sharrar et al. 2020) that perform diverse functions in ecological systems and can mediate symbioses between microbes and plants in certain contexts (Demain and Fang 2000) and ameliorate biotic stress in plants (Ruparelia et al. 2022). Analyzing the biosynthetic gene clusters (BGCs) involved in the production of secondary metabolites within the hazelnut rhizosphere, therefore, provides an opportunity to identify compounds and microbial taxa that can be screened as biological control agents in hazelnut.

In our study, no substantial differences in the distribution of BGC classes were evident between conditions, and all conditions had BGCs encoding the production of non-ribosomal peptides (NRPs), ribosomally synthesized and post-translationally modified peptides (RiPPs), and terpenes. The production of secondary metabolites of these classes has repeatedly been shown to contribute to the biocontrol potential of plant growth-promoting rhizobacteria (PGPRs). For example, *Bacillus* species produce NRPs that can serve as biocontrol agents, such as iturin, surfactin, and fengycin (Ranjan et al. 2023). Likewise, species of the fungal genus *Trichoderma* have demonstrated potential as biocontrol agents (Sood et al. 2020), which could relate to the diversity of terpenoids and complex terpenes produced by these organisms (Bai et al. 2022; Lee et al. 2016).

Many of the MAGs assembled in this study have the genetic potential to produce diverse secondary metabolites. One such example is MAG-30, classified as belonging to the species “*Lysobacter yananisis*”. Species of the genus *Lysobacter* have potential as biocontrol agents (Kilic-Ekici and Yuen 2003; Puopolo et al. 2014), and they produce a range of antibiotics active against numerous plant pathogens (Gómez Expósito et al. 2015; Hayward et al. 2010). Likewise, our *Lysobacter* MAG has the genetic potential to produce various secondary metabolites of relevance for biocontrol, including NRPs, lantipeptides, polyketides, and RiPPs. Another MAG with high biosynthetic potential is MAG-07, classified as belonging to the genus *Actinospica*. Species of this genus have been isolated from diverse ecosystems including forest soils (Kusuma et al. 2022; Cavaletti et al. 2006), and studies suggest that members of this genus have strong biocontrol potential (Qi et al. 2019; Rodriguez-Mena et al. 2022). Indeed, members of the genus *Actinospica* have high genetic potential to produce a wide array of secondary metabolites (Majer et al. 2021). Likewise, our *Actinospica* MAG contains several genomic regions potentially involved in the production of NRPs, lantipeptides, RIPPs, polyketides, and terpenes. Considering these results, it would be interesting to isolate strains of the genera *Lysobacter* and *Actinospica* from hazelnut rhizosphere soils and investigate their potential to serve as biocontrol strains and reduce the susceptibility of hazelnut to *A. anomala* infection.

### Conclusions

The co-evolution of plants and their resident microbial communities has resulted in a complex dynamic where the microbial genetic complement can drastically impact important plant features. Soil microbes participate in important biogeochemical cycles that impact plant nutrient acquisition, while plant-associated microbes can also influence plant immunity. Here, metagenomics was used to examine the microbial communities of three hazelnut rhizosphere sample groups obtained from trees that displayed differing levels of resistance to the pathogen *A. anomala*: *C. americana* (resistant), *C. avellana* (susceptible) and *C. avellana* (resistant). While the rhizosphere samples were differentiated from the bulk soil samples, little taxonomic or functional variation was seen between the different rhizosphere groups. A total of 45 dereplicated, high-quality MAGs were generated, many of which carried multiple BGCs. This included two MAGs representing organisms from the bacterial genera *Actinospica* and *Lysobacter* that have previously been identified as potential biocontrol agents. This study serves to inform future efforts aimed at leveraging hazelnut-associated microbes toward the improvement of diverse hazelnut features including disease outcomes, identifying promising candidate taxa that could be isolated and screened for their capacity to manage *A. anomala* infection. In addition, this study provides insights into the taxonomic diversity and functional potential of the rhizosphere microbial communities of hazelnut, expanding our understanding of the rhizosphere microbiome of woody perennials.

## Data availability

All raw Illumina sequencing data, metagenome assemblies, and MAGs are available from the National Center for Biotechnology Information (NCBI) under BioProject PRJNA1338491 Accessions for individual datasets are provided in **Table S3**. All code to repeat the analyses described in this manuscript is available through GitHub (github.com/diCenzo-Lab/016_2025_hazelnut_rhizosphere_metagenomics).

## ACKNOWLEDGEMENTS

We thank Sharen Roland and other personnel at the Centre d’Expertise et de Services (CES) of Génome Québec for the advice on sequencing strategies and for performing all the Illumina sequencing reported in this study. We would also like to thank Rutgers University Department of Plant Biology for providing access to their facilities for sample collection and preparation. This work was funded by the Natural Sciences and Engineering Research Council of Canada (NSERC) through Discovery Grants to S.R. and G.C.D. and the NSERC Collaborative Research and Training Experience Grant “Genome Editing for Food Security and Environmental Sustainability” to S.R. This work was also supported by the USDA-NIFA Hatch project NJ12171 through the New Jersey Agricultural Experiment Station. This research was enabled, in part, through computational resources provided by Compute Ontario (computeontario.ca) and the Digital Research Alliance of Canada (alliancecan.ca).

