## Supplementary Material for "*Corylus avellana* disease management: using metagenomics to illuminate the rhizosphere microbiome of hazelnut"

**Supplementary Material for**
*Corylus avellana* disease management: using metagenomics to illuminate the rhizosphere microbiome of hazelnut

John K. Robinson, John M. Steele, Thomas J. Molnar, Sharon Regan, George C. diCenzo

George C diCenzo


**This file includes:**

Tables S1 to S3 (pages 2-6)
Figures S1 to S4 (pages 7-10)

**Table S1.** Metagenome sequencing and assembly statistics.

| Sample | Index | Initial read count | Cleaned read count | Metagenome size (bp) | Number of Contigs | N50 (bp) | Total length of contigs ≥ 500bp (bp) | Number of contigs ≥ 500bp | Total length of contigs ≥ 1000bp (bp) | Number of contigs ≥ 1000bp |
|---|---|---|---|---|---|---|---|---|---|---|
| Bulk soil rep. 1 | B2 | 51,880,556 | 50,033,788 | 1,507,285,007 | 3,026,650 | 968,231 | 703,996,213 | 861,863 | 234,194,472 | 143,772 |
| Bulk soil rep.2 | B3 | 56,455,215 | 54,656,811 | 1,760,388,404 | 3,258,114 | 917,834 | 938,534,172 | 1,028,995 | 402,488,425 | 218,639 |
| Bulk soil rep.3 | B4 | 35,138,424 | 31,340,823 | 711,298,515 | 1,508,991 | 513,783 | 295,233,338 | 384,068 | 80,164,916 | 52,453 |
| Bulk soil rep.4 | B5 | 60,972,066 | 58,965,780 | 1,922,313,949 | 3,726,780 | 1,146,156 | 960,548,760 | 1,140,891 | 346,401,406 | 206,742 |
| Native rep.1 | R1P1 | 39,500,132 | 35,283,360 | 704,566,019 | 1,539,916 | 537,985 | 264,886,115 | 347,475 | 69,428,417 | 42,525 |
| Native rep.2 | R1P5 | 51,810,390 | 50,087,267 | 1,473,421,134 | 3,060,954 | 1,024,616 | 637,388,327 | 814,309 | 181,490,030 | 110,838 |
| Native rep.3 | R1P9 | 54,247,765 | 52,536,235 | 1,590,819,716 | 3,282,831 | 1,097,961 | 704,194,531 | 902,341 | 198,277,168 | 124,614 |
| Native rep.4 | R1P14 | 54,787,022 | 52,871,150 | 1,512,961,561 | 3,187,656 | 1,090,529 | 640,062,068 | 843,261 | 163,187,047 | 106,419 |
| Native rep.5 | R1P18 | 40,803,397 | 36,430,522 | 748,213,675 | 1,553,423 | 507,714 | 318,626,581 | 388,261 | 104,162,353 | 58,241 |
| Susceptible rep.1 | R3P3 | 118,478,008 | 113,588,930 | 4,007,941,913 | 7,549,956 | 2,260,483 | 2,097,942,028 | 2,436,581 | 802,311,098 | 472,293 |
| Susceptible rep.2 | R3P5 | 38,658,251 | 34,762,764 | 672,861,461 | 1,485,880 | 524,456 | 245,198,220 | 325,649 | 59,441,879 | 35,604 |
| Susceptible rep.3 | R3P11 | 35,208,735 | 31,398,170 | 544,796,943 | 1,227,732 | 446,052 | 188,460,453 | 260,145 | 38,965,972 | 25,885 |
| Susceptible rep.4 | R3P15 | 56,820,109 | 54,699,006 | 1,502,981,727 | 3,206,815 | 1,111,018 | 614,470,960 | 819,938 | 148,287,978 | 97,022 |
| Susceptible rep.5 | R3P20 | 59,050,822 | 57,205,187 | 1,647,245,580 | 3,512,821 | 1,225,441 | 684,919,232 | 932,723 | 152,022,164 | 108,253 |
| Resistant rep.1 | R3P9- | 49,832,298 | 48,121,647 | 1,278,521,990 | 2,762,123 | 972,500 | 507,187,271 | 687,435 | 116,573,965 | 80,210 |
| Resistant rep.2 | R3P10- | 49,270,339 | 47,503,241 | 1,409,252,362 | 2,906,874 | 966,404 | 621,543,140 | 789,820 | 182,020,828 | 113,495 |
| Resistant rep.3 | R3P12- | 56,900,520 | 55,060,780 | 1,583,954,829 | 3,345,611 | 1,151,098 | 669,932,171 | 891,398 | 167,408,196 | 115,716 |
| Resistant rep.4 | R3P14- | 38,224,297 | 34,180,974 | 668,813,197 | 1,477,114 | 524,236 | 246,298,767 | 332,238 | 56,311,704 | 35,967 |
| Resistant rep.5 | R3P18- | 44,687,280 | 39,740,139 | 806,693,915 | 1,807,203 | 658,640 | 288,787,441 | 406,042 | 54,149,209 | 39,267 |

**Table S2.** Statistics of taxonomic classification of Illumina reads using Kraken2 or MMSeqs2.

| Sample | Index | Reads classified by Kraken2 (%) | Reads classified based on MMSeqs2 output (%) * | Reads classified as bacteria based on MMSeqs2 output (%) | | Reads classified as archaea based on MMSeqs2 output (%) | | Reads classified as eukaryotes based on MMSeqs2 output (%) | |
|---|---|---|---|---|---|---|---|---|---|
| | | | | Of all reads | Of classified reads | Of all reads | Of classified reads | Of all reads | Of classified reads |
| Bulk soil rep. 1 | B2 | 0.8 | 19.47 | 18.3 | 94 | 0.74 | 3.8 | 0.11 | 0.56 |
| Bulk soil rep.2 | B3 | 1.04 | 31.18 | 29.33 | 94.07 | 1.22 | 3.91 | 0.11 | 0.35 |
| Bulk soil rep.3 | B4 | 1.03 | 11.12 | 10.02 | 90.11 | 0.71 | 6.38 | 0.06 | 0.54 |
| Bulk soil rep.4 | B5 | 1.77 | 24.61 | 23.56 | 95.73 | 0.24 | 0.97 | 0.23 | 0.93 |
| Native rep.1 | R1P1 | 1.44 | 8.08 | 7.18 | 88.86 | 0.31 | 3.84 | 0.35 | 4.33 |
| Native rep.2 | R1P5 | 1.47 | 16.41 | 15.46 | 94.21 | 0.36 | 2.19 | 0.19 | 1.16 |
| Native rep.3 | R1P9 | 1.39 | 16.76 | 15.58 | 92.96 | 0.56 | 3.34 | 0.19 | 1.13 |
| Native rep.4 | R1P14 | 1.35 | 15.2 | 14.42 | 94.87 | 0.24 | 1.58 | 0.16 | 1.05 |
| Native rep.5 | R1P18 | 0.95 | 11.38 | 10.38 | 91.21 | 0.48 | 4.22 | 0.23 | 2.02 |
| Susceptible rep.1 | R3P3 | 1.07 | 27.09 | 25.07 | 92.54 | 0.39 | 1.44 | 1.04 | 3.84 |
| Susceptible rep.2 | R3P5 | 1.29 | 7.71 | 6.39 | 82.88 | 0.4 | 5.19 | 0.1 | 1.3 |
| Susceptible rep.3 | R3P11 | 1.26 | 5.99 | 5.17 | 86.31 | 0.57 | 9.52 | 0.05 | 0.83 |
| Susceptible rep.4 | R3P15 | 1.39 | 13.99 | 13.33 | 95.28 | 0.19 | 1.36 | 0.17 | 1.21 |
| Susceptible rep.5 | R3P20 | 1.4 | 14.2 | 13.49 | 95 | 0.25 | 1.76 | 0.14 | 0.99 |
| Resistant rep.1 | R3P9- | 1.21 | 12.24 | 11.59 | 94.69 | 0.17 | 1.39 | 0.14 | 1.14 |
| Resistant rep.2 | R3P10- | 1.32 | 16.09 | 15.05 | 93.54 | 0.37 | 2.3 | 0.25 | 1.55 |
| Resistant rep.3 | R3P12- | 1.21 | 14.41 | 13.57 | 94.17 | 0.35 | 2.43 | 0.17 | 1.18 |
| Resistant rep.4 | R3P14- | 1.24 | 8.49 | 7.81 | 91.99 | 0.35 | 4.12 | 0.12 | 1.41 |
| Resistant rep.5 | R3P18- | 1.13 | 7.11 | 6.29 | 88.47 | 0.53 | 7.45 | 0.09 | 1.27 |

* Assembled contigs of at least 500 bp were taxonomically classified using MMSeqs2, and the taxonomic classifications were then propagated to the Illumina reads that mapped to each contig.

**Table S3.** Taxonomic classification and genome statistics for the metagenome-assembled genomes (MAGs) produced in this study.

| MAG name | GTDB-Tk classification * | Genome size (bp) | Number of contigs | Completeness (%) † | Contamination (%) † |
|---|---|---|---|---|---|
| Archaea | | | | | |
| Nitrososphaeraceae_archaeon_QU-JR-MAG-02 | p_Thermoproteota; c_Nitrososphaeria; o_Nitrososphaerales; f_Nitrososphaeraceae; g_JAFAQB01; s_ | 3,003,512 | 1,180 | 83.82 | 8.08 |
| Nitrososphaeraceae_archaeon_QU-JR-MAG-09 | p_Thermoproteota; c_Nitrososphaeria; o_Nitrososphaerales; f_Nitrososphaeraceae; g_JAFAQB01; s_ | 3,830,128 | 1,261 | 86.96 | 4.37 |
| Nitrososphaeraceae_archaeon_QU-JR-MAG-21 | p_Thermoproteota; c_Nitrososphaeria; o_Nitrososphaerales; f_Nitrososphaeraceae; g_JARBAU01; s_ | 2,739,335 | 439 | 97.25 | 1.46 |
| Nitrososphaeraceae_archaeon_QU-JR-MAG-33 | p_Thermoproteota; c_Nitrososphaeria; o_Nitrososphaerales; f_Nitrososphaeraceae; g_TA-21; s_TA-21 sp021324715 | 1,786,828 | 223 | 95.79 | 0.97 |
| Bacteria | | | | | |
| Vicinamibacterales_bacterium_QU-JR-MAG-01 | p_Acidobacteriota; c_Vicinamibacteria; o_Vicinamibacterales; f_2-12-FULL-66-21; g_JALZOC01; s_ | 7,761,909 | 362 | 97.25 | 6.84 |
| Solirubrobacterales_bacterium_QU-JR-MAG-03 | p_Actinomycetota; c_Thermoleophilia; o_Solirubrobacterales; f_70-9; g_VAYN01; s_ | 2,633,149 | 123 | 96.98 | 0.95 |
| Solirubrobacterales_bacterium_QU-JR-MAG-04 | p_Actinomycetota; c_Thermoleophilia; o_Solirubrobacterales; f_70-9; g_VRUE01; s_ | 2,723,173 | 130 | 94.68 | 2.16 |
| Thermoanaerobaculia_bacterium_QU-JR-MAG-05 | p_Acidobacteriota; c_Thermoanaerobaculia; o_Gp7-AA8; f_Gp7-AA8; g_QHVT01; s_ | 6,747,948 | 486 | 97.10 | 5.24 |
| Gaiellales_bacterium_QU-JR-MAG-06 | p_Actinomycetota; c_Thermoleophilia; o_Gaiellales; f_JAICJC01; g_JAICYJ01; s_ | 3,264,548 | 833 | 86.28 | 6.48 |
| Actinospica_sp_QU-JR-MAG-07 | p_Actinomycetota; c_Actinomycetes; o_Streptomycetales; f_Catenulisporaceae; g_Actinospica; s_ | 7,842,952 | 1,423 | 91.30 | 7.89 |
| Pseudolabrys_sp_QU-JR-MAG-08 | p_Pseudomonadota; c_Alphaproteobacteria; o_Rhizobiales; f_Xanthobacteraceae; g_Pseudolabrys; s_ | 3,767,974 | 849 | 82.28 | 2.24 |
| Rokubacteriales_bacterium_QU-JR-MAG-10 | p_Methylomirabilota; c_Methylomirabilia; o_Rokubacteriales; f_CSP1-6; g_AR5; s_ | 6,503,514 | 1,216 | 77.72 | 9.36 |
| Hyphomicrobiaceae_bacterium_QU-JR-MAG-11 | p_Pseudomonadota; c_Alphaproteobacteria; o_Rhizobiales; f_Hyphomicrobiaceae; g_AWTP1-13; s_ | 9,461,451 | 875 | 96.67 | 7.30 |
| Thermoanaerobaculia_bacterium_QU-JR-MAG-12 | p_Acidobacteriota; c_Thermoanaerobaculia; o_Gp7-AA8; f_Gp7-AA8; g_; s_ | 5,041,695 | 1,072 | 87.61 | 3.76 |
| Methyloceanibacter_sp_QU-JR-MAG-13 | p_Pseudomonadota; c_Alphaproteobacteria; o_Rhizobiales; f_Methyloligellaceae; g_Methyloceanibacter; s_ | 1,760,970 | 358 | 72.75 | 1.41 |
| Nitrospira_bacterium_QU-JR-MAG-14 | p_Nitrospirota; c_Nitrospiria; o_Nitrospirales; f_Nitrospiraceae; g_Nitrospira_C; s_Nitrospira_C sp025935475 | 2,940,148 | 1,013 | 87.02 | 5.41 |
| Multivoradales_bacterium_QU-JR-MAG-15 | p_Acidobacteriota; c_Thermoanaerobaculia; o_Multivoradales; f_UBA5704; g_UBA5704; s_ | 5,509,139 | 1,026 | 92.05 | 1.93 |
| Binatia_bacterium_QU-JR-MAG-16 | p_Desulfobacterota_B; c_Binatia; o_UBA9968; f_UBA9968; g_DP-1; s_ | 7,032,466 | 1,862 | 84.90 | 4.23 |
| Limnocylindrales_bacterium_QU-JR-MAG-17 | p_Chloroflexota; c_Limnocylindria; o_Limnocylindrales; f_CSP1-4; g_SPCO01; s_ | 2,998,850 | 781 | 87.66 | 5.10 |
| Solirubrobacterales_bacterium_QU-JR-MAG-18 | p_Actinomycetota; c_Thermoleophilia; o_Solirubrobacterales; f_70-9; g_AC-56; s_ | 2,230,084 | 730 | 79.02 | 6.82 |
| Vicinamibacterales_bacterium_QU-JR-MAG-19 | p_Acidobacteriota; c_Vicinamibacteria; o_Vicinamibacterales; f_UBA2999; g_JAENWD01; s_ | 4,729,133 | 1,637 | 70.01 | 6.86 |
| Marmoricola_sp_QU-JR-MAG-20 | p_Actinomycetota; c_Actinomycetes; o_Propionibacteriales; f_Nocardioidaceae; g_Marmoricola; s_ | 3,438,173 | 1,226 | 74.78 | 8.12 |
| Sphingomicrobium_sp_QU-JR-MAG-22 | p_Pseudomonadota; c_Alphaproteobacteria; o_Sphingomonadales; f_Sphingomonadaceae; g_Sphingomicrobium; s_ | 2,879,127 | 276 | 98.15 | 2.08 |
| Beijerinckiaceae_bacterium_QU-JR-MAG-23 | p_Pseudomonadota; c_Alphaproteobacteria; o_Rhizobiales; f_Beijerinckiaceae; g_JAFASC01; s_ | 4,275,855 | 1,363 | 79.50 | 3.96 |
| Methyloceanibacter_sp_QU-JR-MAG-24 | p_Pseudomonadota; c_Alphaproteobacteria; o_Rhizobiales; f_Methyloligellaceae; g_Methyloceanibacter; s_ | 2,418,542 | 252 | 88.02 | 1.87 |
| Gaiellaceae_bacterium_QU-JR-MAG-25 | p_Actinomycetota; c_Thermoleophilia; o_Gaiellales; f_Gaiellaceae; g_; s_ | 2,184,336 | 656 | 71.09 | 9.64 |
| Solirubrobacterales_bacterium_QU-JR-MAG-26 | p_Actinomycetota; c_Thermoleophilia; o_Solirubrobacterales; f_70-9; g_VRUE01; s_ | 2,405,082 | 209 | 91.24 | 4.60 |
| Pseudolabrys_sp_QU-JR-MAG-27 | p_Pseudomonadota; c_Alphaproteobacteria; o_Rhizobiales; f_Xanthobacteraceae; g_Pseudolabrys; s_ | 3,880,196 | 232 | 95.37 | 0.63 |

| | | | | | |
|---|---|---|---|---|---|
| Pyrinomonadaceae_bacterium_QU-JR-MAG-28 | p_Acidobacteriota; c_Blastocatellia; o_Pyrinomonadales; f_Pyrinomonadaceae; g_UBA11740; s_ | 3,728,135 | 1,181 | 73.69 | 4.20 |
| Streptosporangiaceae_bacterium_QU-JR-MAG-29 | p_Actinomycetota; c_Actinomycetes; o_Streptosporangiales; f_Streptosporangiaceae; g_Palsa-504; s_ | 8,130,091 | 2,214 | 83.59 | 8.47 |
| Lysobacter_yananisis_MAG30 | p_Pseudomonadota; c_Gammaproteobacteria; o_Xanthomonadales; f_Xanthomonadaceae; g_Lysobacter; s_Lysobacter yananisis | 4,410,919 | 1,419 | 76.30 | 4.20 |
| Solirubrobacterales_bacterium_QU-JR-MAG-31 | p_Actinomycetota; c_Thermoleophilia; o_Solirubrobacterales; f_70-9; g_VAYN01; s_ | 2,329,140 | 292 | 76.67 | 3.02 |
| Gaiellaceae_bacterium_QU-JR-MAG-32 | p_Actinomycetota; c_Thermoleophilia; o_Gaiellales; f_Gaiellaceae; g_Palsa-739; s_ | 2,997,075 | 424 | 86.05 | 4.74 |
| Gaiellaceae_bacterium_QU-JR-MAG-34 | p_Actinomycetota; c_Thermoleophilia; o_Gaiellales; f_Gaiellaceae; g_Palsa-739; s_ | 2,810,457 | 294 | 93.95 | 8.48 |
| Solirubrobacterales_bacterium_QU-JR-MAG-35 | p_Actinomycetota; c_Thermoleophilia; o_Solirubrobacterales; f_70-9; g_VAYN01; s_ | 2,589,232 | 76 | 95.98 | 0.57 |
| Gemmatimonadales_bacterium_QU-JR-MAG-36 | p_Gemmatimonadota; c_Gemmatimonadetes; o_Gemmatimonadales; f_GWC2-71-9; g_JACDDX01; s_ | 3,974,458 | 560 | 79.61 | 2.26 |
| Solirubrobacterales_bacterium_QU-JR-MAG-37 | p_Actinomycetota; c_Thermoleophilia; o_Solirubrobacterales; f_70-9; g_VRUE01; s_ | 2,512,822 | 97 | 92.41 | 6.59 |
| Ilumatobacteraceae_bacterium_QU-JR-MAG-38 | p_Actinomycetota; c_Acidimicrobiia; o_Acidimicrobiales; f_Ilumatobacteraceae; g_; s_ | 3,561,652 | 1,268 | 73.91 | 4.70 |
| Gaiellaceae_bacterium_QU-JR-MAG-39 | p_Actinomycetota; c_Thermoleophilia; o_Gaiellales; f_Gaiellaceae; g_; s_ | 2,631,093 | 719 | 70.20 | 4.29 |
| Rhizobium_sp_QU-JR-MAG-40 | p_Pseudomonadota; c_Alphaproteobacteria; o_Rhizobiales; f_Rhizobiaceae; g_Rhizobium; s_ | 5,351,430 | 704 | 94.70 | 2.58 |
| Solirubrobacterales_bacterium_QU-JR-MAG-41 | p_Actinomycetota; c_Thermoleophilia; o_Solirubrobacterales; f_70-9; g_VAYN01; s_ | 1,922,208 | 411 | 84.89 | 5.25 |
| Udaeobacter_sp_QU-JR-MAG-42 | p_Verrucomicrobiota; c_Verrucomicrobiae; o_Chthoniobacterales; f_UBA10450; g_Udaeobacter; s_ | 3,248,333 | 323 | 94.93 | 4.77 |
| Mycobacteriales_bacterium_QU-JR-MAG-43 | p_Actinomycetota; c_Actinomycetes; o_Mycobacteriales; f_CADCTP01; g_CADCTP01; s_ | 4,911,930 | 1,207 | 84.51 | 3.70 |
| Actinomycetota_bacterium_QU-JR-MAG-44 | p_Actinomycetota; c_UBA4738; o_UBA4738; f_HRBIN12; g_JAJNBA01; s_ | 1,840,927 | 374 | 73.35 | 2.56 |
| Gaiellaceae_bacterium_QU-JR-MAG-45 | p_Actinomycetota; c_Thermoleophilia; o_Gaiellales; f_Gaiellaceae; g_GMQP-bins7; s_ | 2,474,174 | 266 | 94.74 | 2.67 |

* Taxonomy determined using the Genome Taxonomy Database Toolkit (GTDB-Tk) with the R220 database. Prefixes designate phylum (p_), class (c_), order (o_), family (f_), genus (g_), and species (s_).

† Genome completeness and contamination were determined using CheckM.

**Table S4.** NCBI accessions to access the genomic data generated in this study. At the time of submission, metagenome assemblies and MAGs had not yet been processed; this table will be updated with accessions for the metagenome assemblies and MAGs once available.

| **Illumina read sets** | |
|---|---|
| **Sample** | **SRA accession** |
| Bulk soil rep. 1 | SRR35753086 |
| Bulk soil rep.2 | SRR35753085 |
| Bulk soil rep.3 | SRR35753075 |
| Bulk soil rep.4 | SRR35753074 |
| Native rep.1 | SRR35753073 |
| Native rep.2 | SRR35753072 |
| Native rep.3 | SRR35753071 |
| Native rep.4 | SRR35753070 |
| Native rep.5 | SRR35753069 |
| Susceptible rep.1 | SRR35753068 |
| Susceptible rep.2 | SRR35753084 |
| Susceptible rep.3 | SRR35753083 |
| Susceptible rep.4 | SRR35753082 |
| Susceptible rep.5 | SRR35753081 |
| Resistant rep.1 | SRR35753080 |
| Resistant rep.2 | SRR35753079 |
| Resistant rep.3 | SRR35753078 |
| Resistant rep.4 | SRR35753077 |
| Resistant rep.5 | SRR35753076 |

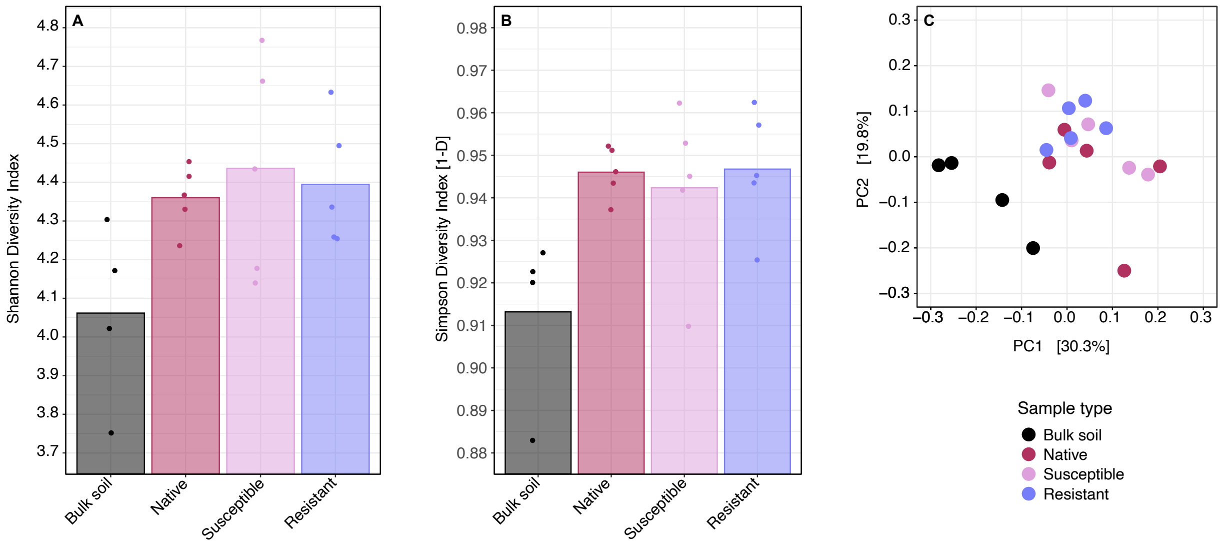


**Figure S1. Diversity metrics of subsampled hazelnut rhizosphere samples.** The Illumina read sets were subsampled such that there was an equal number of reads used for metagenome assembly for each sample. Diversity statistics were calculated following the down-sampling. (**A**, **B**) Alpha diversity values calculated using the (A) Shannon Diversity Index (Kruskal-Wallis rank sum test, p-value = 0.12) or (B) Simpson Diversity Index [1-D] (Kruskal-Wallis rank sum test, p-value = 0.07) are shown. Alpha diversity was calculated based on species level taxonomic classification of assembled contigs (minimum length of 500 bp). (**C**) A Principal Coordinate Analysis (PCoA) plot based on Bray-Curtis dissimilarity values is shown (PERMANOVA, p-value < 0.001), with pairwise adonis tests indicating that the bulk soil was statistically different from the rhizospheres of the native (p.adjusted = 0.047), susceptible (p.adjusted = 0.048), and resistant (p.adjusted = 0.048) varieties.

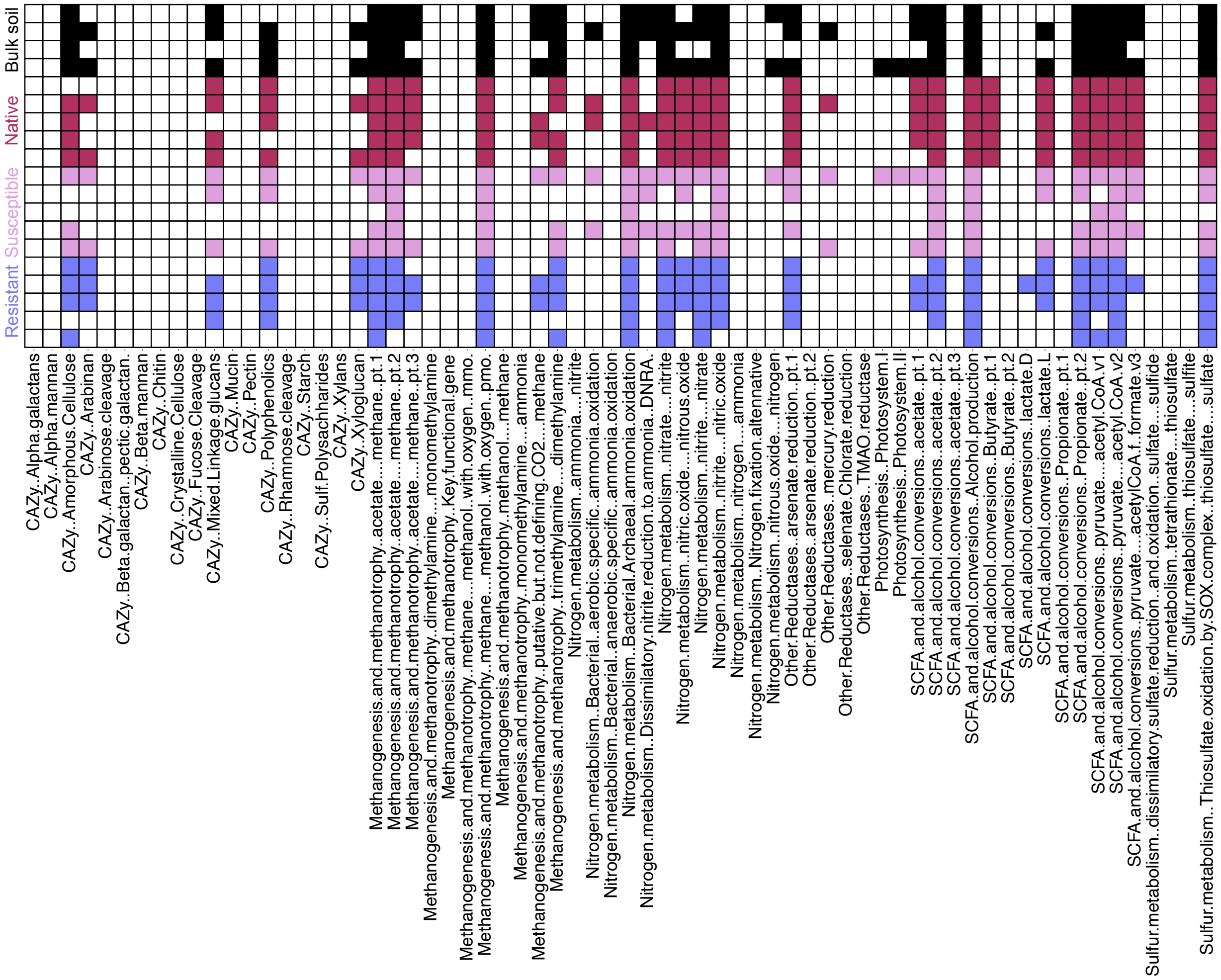


**Figure S2. Presence or absence of various metabolic pathways across metagenomic samples.** DRAM was used to functionally annotate each of the metagenomes, and the output used to determine the presence or absence of various metabolic pathways related to carbohydrate metabolism, methanogenesis and methanotrophy, nitrogen metabolism, short-chain fatty acid and alcohol conversions, and sulfur metabolism. The rows represent each of the metagenomic samples, colour coded by group, while the columns represent different metabolic pathways. A coloured box indicates that the pathway was detected in a given metagenome, whereas a white box indicates that the pathway was not detected.

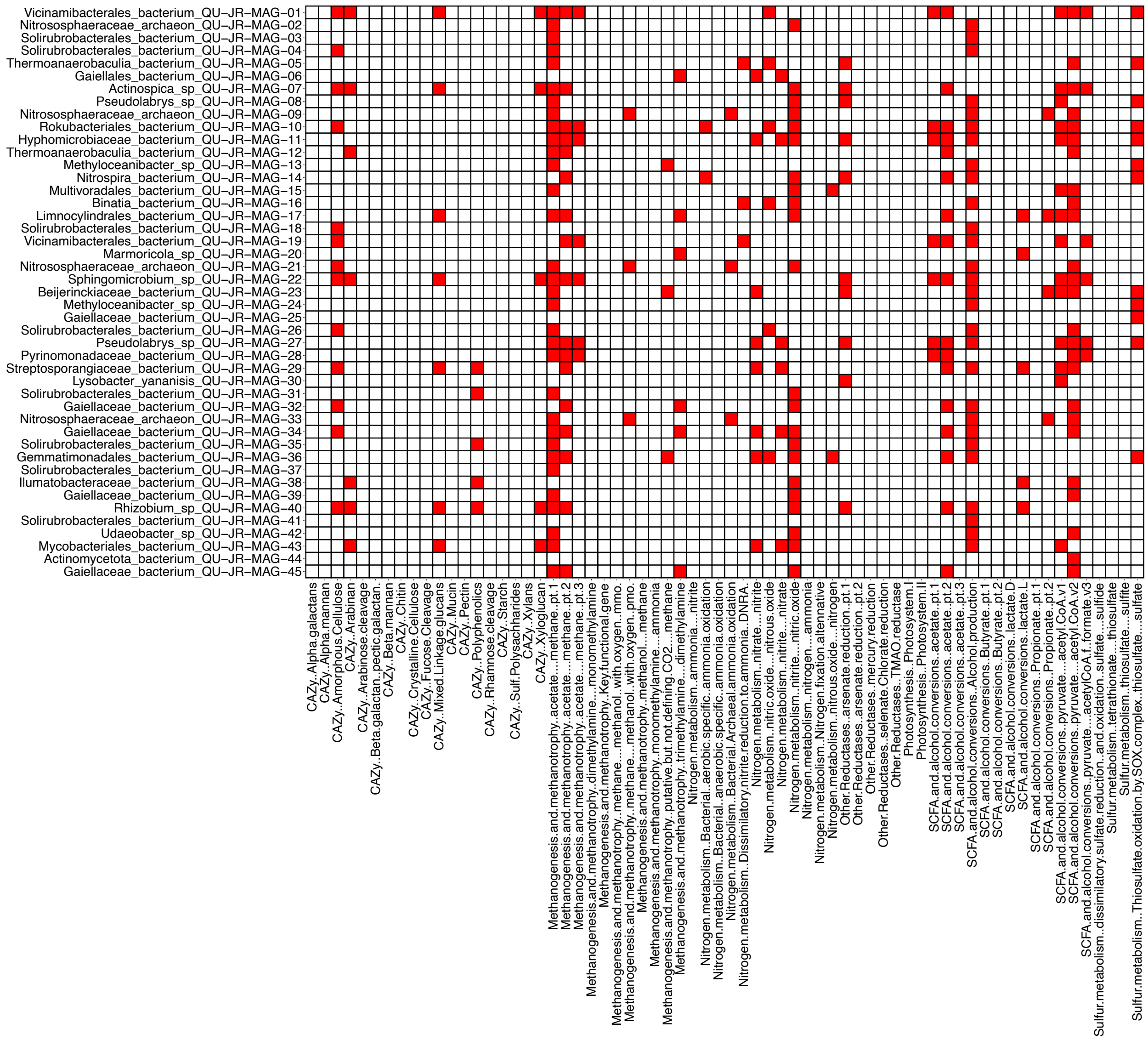


**Figure S3. Presence or absence of various metabolic pathways across MAGs.** DRAM was used to functionally annotate each of the metagenome-assembled genomes (MAGs), and the output used to determine the presence or absence of various metabolic pathways related to carbohydrate metabolism, methanogenesis and methanotrophy, nitrogen metabolism, short-chain fatty acid and alcohol conversions, and sulfur metabolism. The rows represent each of the MAGs, while the columns represent different metabolic pathways. A red box indicates that the pathway was detected in a given MAG, whereas a white box indicates that the pathway was not detected.

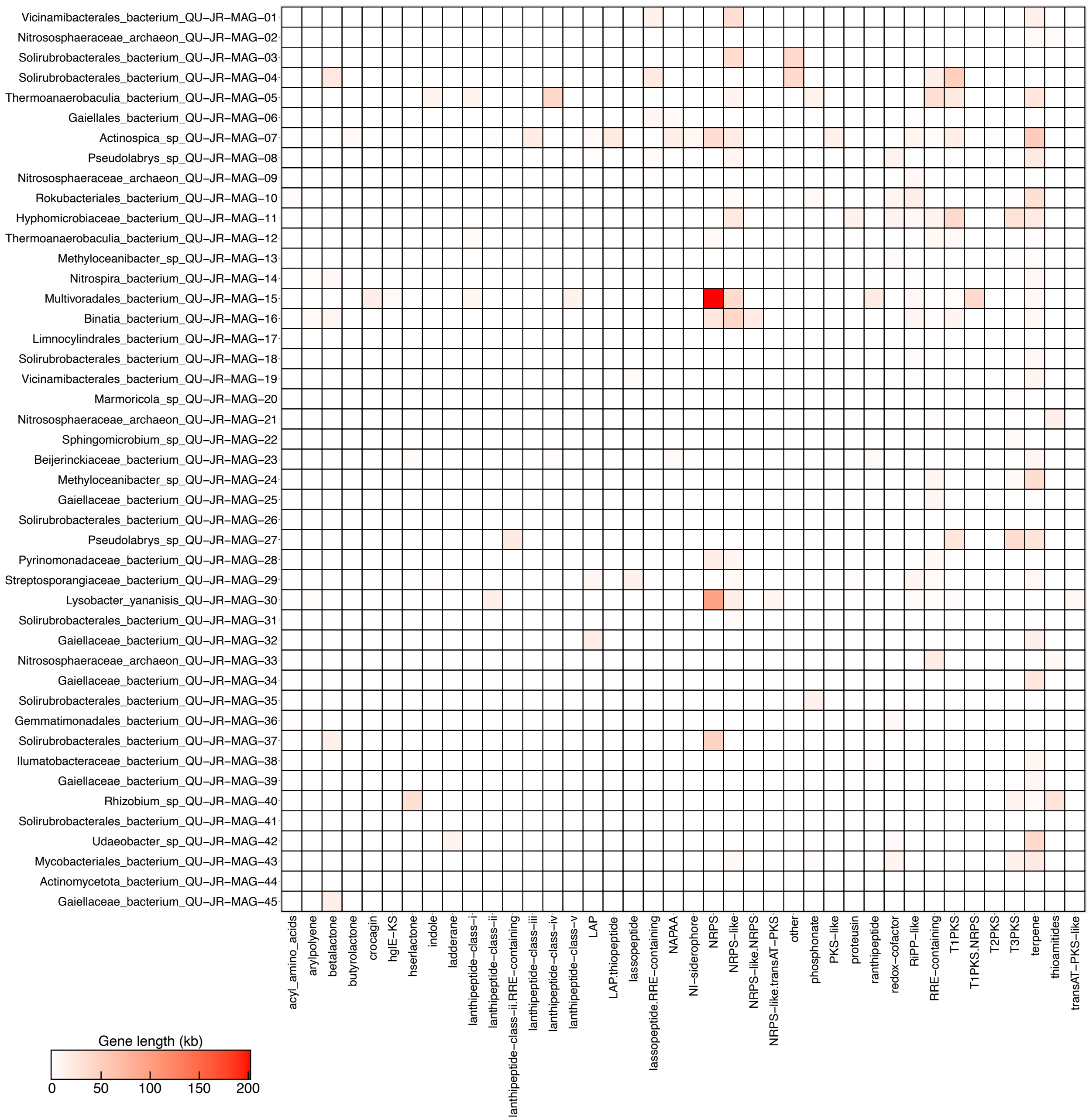


**Figure S4. Putative biosynthetic gene clusters detected in the MAGs.** A heatmap showing the cumulative length of predicted biosynthetic gene clusters summarized by predicted product (columns) in each of the metagenome-assembled genomes (MAGs) (rows). The colour of a box represents the cumulative length (kb) of biosynthetic gene clusters detected that encode a predicted product, with deeper red indicating higher counts.
